# Artificial Intelligence Resolves Kinetic Pathways of Magnesium Binding to RNA

**DOI:** 10.1101/2021.07.25.453696

**Authors:** Jan Neumann, Nadine Schwierz

## Abstract

Magnesium is an indispensable cofactor in countless vital processes. In order to understand its functional role, the characterization of the binding pathways to biomolecules such as RNA is crucial. Despite the importance, a molecular description is still lacking since the transition from the water-mediated outer-sphere to the direct inner-sphere conformation is on the millisecond timescale and therefore out of reach for conventional simulation techniques. To fill this gap, we use transition path sampling to resolve the binding pathways and to elucidate the role of the solvent in the reaction. The results reveal that the molecular void provoked by the leaving phosphate oxygen of the RNA is immediately filled by an entering water molecule. In addition, water molecules from the first and second hydration shell couple to the concerted exchange. To capture the intimate solute-solvent coupling, we perform a committor analysis as basis for a machine learning algorithm that derives the optimal deep learning model from thousands of scanned architectures using hyperparameter tuning. The results reveal that the properly optimized deep network architecture recognizes the important solvent structures, extracts the relevant information and predicts the commitment probability with high accuracy. Our results provide a quantitative description of solute-solvent coupling which is ubiquitous for kosmotropic ions and governs a large variety of biochemical reactions in aqueous solutions.

## INTRODUCTION

Magnesium plays a vital role in almost every biological process. By now more than 800 different biochemical roles of Mg^2+^ have been identified in physiological possesses ranging from the creation of cellular energy or the synthesis of biomolecules to the activation of enzymes and ribozymes [1–5]. The specific requirement for Mg^2+^ as a cofactor is particularly pronounced in nucleic acid systems where Mg^2+^ plays structural roles by complexing negatively charged groups or catalytic roles by accelerating or inhibiting chemical reactions in ribozymes [3, 6–8]. Despite the importance, the molecular pathways of Mg^2+^ binding to RNA have not been resolved to date and the dynamic interplay of direct ion-RNA and indirect water-mediated interactions remains elusive.

In aqueous solutions, the first hydration shell of Mg^2+^ consists of six water molecules arranged in octahedral symmetry [9]. Water molecules from the first hydration shell exchange with the second, more loosely bound hydration shell on the microsecond timescale [10–12]. This dynamic equilibrium allows for ligand exchange. Hereby, oxygen atoms are the preferred binding partners, in particular the non-bridging oxygens of the phosphate group on the backbone of RNA [3, 13]. Interestingly, water exchange around Mg^2+^ is orders of magnitude slower compared to other metal ions [10, 14]. The long lifetimes of water molecules in the first hydration shell facilitate two distinct binding arrangements: Inner-sphere and outer-sphere. In inner-sphere binding, one water molecule is removed from the first hydration shell and the ion is in direct contact with the RNA functionalities. In outer-sphere binding, the interactions with the RNA are mediated by water while Mg^2+^ remains coordinated by six water molecules [3].

Does Mg^2+^ bind in inner- or outer-sphere conformation and how can it transition from the one binding mode to the other? Discerning the nature of the exact interactions is an important and controversially discussed question in the field [13, 15, 16]. Typically, structural knowledge on the binding mode of Mg^2+^ is obtained from crystallographic experiments. However, the correct assignment in the electron density maps remains notoriously difficult. Yet, with proper stereochemical guidelines a consistent picture emerges: Nucleobase nitrogen and carbonyls are poor inner-sphere Mg^2+^ binders [15, 16] while the non-bridging oxygens of the phosphate group are the primary nucleic acid binding location. In large RNA structures, Mg^2+^ ions can, depending on the exact environment, bind in inner- or outer-sphere conformation [3, 17] whereas the inner-sphere conformation is thermodynamically favored in simple mono-nucleotide systems [18].

In addition to the structural information from crystallography, NMR can provide valuable insight into the exchange kinetics [10, 19, 20]. Depending on the number of direct Mg^2+^-RNA contacts, the timescale for ligand exchange ranges from milliseconds [19, 21, 22] to hundreds of seconds [20]. However, the structural changes at the microscopic level during an exchange process are not accessible from experiments. Here, simulations can contribute important insights by characterizing the solvent behavior and by providing a unique atomistic description of the dynamics. Yet, simulating the transition from outer-to-inner sphere binding is tremendously challenging for two reasons: According to experiments, the transition is on the millisecond to second timescale and therefore out of reach for conventional all-atom simulations. Alternatively, enhanced sampling techniques such as metadynamics, replica exchange and umbrella sampling can be applied [23, 24] but do not prove any information on the exchange dynamics and reaction mechanism.

The second challenge is the coupling of the solvent to the exchange dynamics. Kosmotropic ions, such as Mg^2+^, provoke strong structural ordering of the first hydration shells. Therefore, any reaction in aqueous solutions is expected to be governed by a complex interplay of structural, orientational and hydrogen bonding effects extending over several hydration shells. It is therefore not surprising that the solvent is not only a spectator of the chemical process but plays an active role in the evolution of the reaction. Similarly, the dynamics of seemingly simple processes such as dipeptide isomerization, ion pair formation, water exchange between hydration shells, or proton and electron transfer is governed by solute-solvent coupling [25–33]. Therefore, attempts in describing the dynamics of such processes in terms of simplified reaction coordinates that do not include the solute-solvent coupling are likely to fail for several reasons. (i) Solvent reorganization is orthogonal to such simplified reaction coordinates giving rise to pronounced non-Markovian behavior or memory effects [34]. (ii) The metastable states are not uniquely separated leading to the violation of the no-recrossing assumption and failure of transition state theory [35]. (iii) Enhanced sampling techniques that rely on biasing the slow degrees of freedom become inefficient as the solvent reorganization becomes slower than the reactive motion itself. Consequently, quantifying the contribution of the solvent to the reaction coordinate is one of the long-standing problems in chemical reaction kinetics.

In order to address both challenges, we apply transition path sampling as a particularly powerful sampling strategy to provide unbiased microscopic insight into the dynamics of Mg^2+^ binding to the phosphate oxygen of RNA. Subsequently, we perform a committor analysis comprising more than 28,600 conformations as basis for a machine learning algorithm which automatically selects the optimal deep learning model from thousands of scanned architectures in a robust and efficient manner. The resulting optimized deep neural network is shown to capture the intimate solute-solvent coupling and to provide an accurate description of the complete dynamical process.

## METHODS

### Atomistic model and simulation setup

Our model system consists of an RNA dinucleotide with two guanine nucleobases. We investigate Mg^2+^ association at the non-bridging phosphate oxygens since they are the primary nucleic acid binding site for Mg^2+^ [3, 16]. Since both oxygen atoms (O1P and O2P) have identical force field parameters, we focus on the interactions with O1P which has a partial charge of −0.776*e*. A single Mg^2+^ ion, one Cl^−^ ion, and 2150 water molecules are added to the cubic simulation box (L = 40 Å). For Mg^2+^ and Cl^−^, recently optimized force field parameters were used [36] in combination with the TIP3P water model [37]. The TIP3P water model assigns partial charges of −0.834*e* and 0.417*e* to oxygen and hydrogen. The force field parameters of the RNA are taken from Amber99sb-ildn* [38] with parmbsc0 [39] and *χ*_0*L*3_ corrections [40]. The TIP3P water model was used since the RNA force field has been optimized in combination with TIP3P water. The Mg^2+^ parameters were optimized in our previous work and are particularly suited to investigate ion binding since they reproduce experimental activity coefficients, the hydration free energy and the interchange dissociative water exchange mechanism [36, 41].

All simulations were performed using GROMACS [42] with periodic boundary conditions. Particle mesh Ewald summation was used and a Fourier spacing of 0.12 nm and a grid interpolation up to order 4 to handle long-range electrostatic forces. Close Coulomb real space interactions were cut off at 1.2 nm and Lennard-Jones (LJ) interactions after 1.2 nm, respectively. Long-range dispersion corrections for energy and pressure were applied to account for errors stemming from truncated LJ interactions.

The initial energy minimization was performed with the steepest descent algorithm. For each simulation, a NVT and a subsequent NPT equilibration was done for 1 ns, controlling the temperature at 300 K and at a pressure of 1 bar with the Berendsen thermostat and barostat [43]. All production runs and the transition path sampling were done in the NVT ensemble at a temperature of 300 K using the velocity rescaling thermostat with stochastic term [44] and a time step of 2 fs. Here, the velocity rescaling thermostat was used since it generates the canonical ensemble and ensures detailed balance in the canonical ensemble of transition paths [45].

### Free energy profiles

The one-dimensional free energy profile as a function of the distance *r*_*I*_ between the Mg^2+^ ion and the oxygen atom O1P were calculated from umbrella sampling using PLUMED [46]. A force constant *k*_*b*_ = 600, 000 kJ/(mol nm^2^) and a window spacing of 0.005 nm was used for *r*_*I*_ < 0.35 nm. A force constant *k*_*b*_ = 60, 000 kJ/(mol nm^2^) and a window spacing of 0.01 nm was used for *r*_*I*_ > 0.35 nm.

The two-dimensional free energy profiles as a function of *r*_*I*_ and the effective hydration distance *s*_6_ were calculated from umbrella sampling using PLUMED [46]. Hereby, *s*_6_ was defined as the sum of six Mg^2+^-oxygen distances

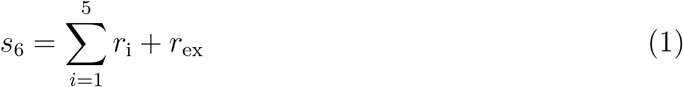

where *r*_i_ are the distances between Mg^2+^ and five closest water molecules and *r*_ex_ is the distance between Mg^2+^ and the exchanging water molecule. A force constant *k*_*b*_ = 100, 000 kJ/(mol nm^2^) and a window spacing of 0.01 nm was used. Without further restraints, the system shows three stable states in the two-dimensional projection (see Figure S1 in the Supporting Information). Without loss of generality, we focus only on transitions between the two relevant states (corresponding to the inner-sphere conformation with 5 coordinating water molecules and the outer-sphere conformation in which O1P is replaced by the selected water molecule for which the umbrella potential is applied). This is achieved by an additional biasing potential on the hydration number (see Supporting Information for further details). Position restraints were applied on all heavy atoms of the dinucleotide with exception of the non-bridging oxygen atoms of the phosphate group. Each umbrella simulation was performed for 2 ns discarding 500 ps for equilibration. The free energy profiles were calculated using the weighted histogram analysis method (WHAM) [47].

### Transition state theory

TST is the most popular theory to calculate reaction rates. In simple systems for which the reaction coordinate is exactly known, TST gives an accurate estimate of the rate. However, in complex many body systems as the one presented here, TST could fail due to the violation of the non-recrossing hypothesis which forms the cornerstone of the theory. Therefore, TST can be used only to provide an upper estimated for the rate constant. For a more accurate estimate, additional corrections as implemented in the reactive flux method are required [35, 48] but are beyond the scope of our current work. Following conventional transition state theory (TST), the upper estimate of the rate constant is calculated from [49, 50]

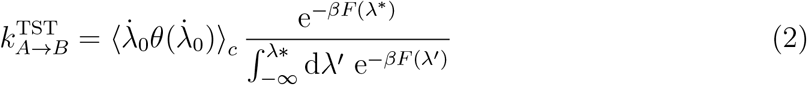

where *λ*\* is the position of the barrier top, 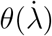 is the Heaviside step function and ⟨…⟩_*c*_ denotes the average over the restrained ensemble of trajectories initiated from an equilibrium ensemble of phase points on the dividing surface.

### Transition path sampling

Transition path sampling [45, 51] was used to harvest an ensemble of rare trajectories that connect the two stable states. Starting from an initial reactive pathway generated at high temperature, new trial trajectories were created by randomly selecting a time slice, randomizing the velocities and integrating the equations of motion forward and backward in time. Using standard two-way shooting moves with a fixed length of 1.6 ps, trial moves are accepted if they connected the two stable states and rejected otherwise.

### Committor analysis and transition states

For a simulation snapshot taken from a reactive transition pathway, the committor *p*_A_ is defined as the probability of the conformation initiated with randomized velocities drawn from a Maxwell-Boltzmann distribution to be committed to state A (inner-sphere conformation). Conformations from basin A have *p*_A_ = 1, conformations from basin B have *p*_A_ = 0 and transition states have *p*_A_ = 0.5. The commitment probability *p*_A_ was calculated from the fraction of trajectories initiated with randomized velocities that reach state A. 29,112 conformations along more than 2500 independent pathways obtained from transition path sampling were used as shooting points. 28,612 shooting points were chosen in the transition region (with 0.6 < *λ* < 1.0 based on the coordinate *λ* defined in Eq. 7). In addition, 500 shooting points were selected in the regions of the stable states (*λ* < 0.6 and *λ* > 1.0). For each shooting point, 100 trajectories were initiated with random velocities and run forward and backward for 2 ps. A conformation was identified as transition state if half of the trajectories relaxed into each stable state (0.45 < *p*_A_ < 0.55). From the ensemble of shooting point, two subsets with *r*_*I*_ = 0.28 ± 0.02 or *λ* = 0.8 ± 0.05 with 3,500 randomly drawn datapoints were selected and the distribution *p*(*p*_A_) was calculated.

### Commitment probability from deep neural networks

Deducing an appropriate description by visual inspection is virtually impossible if complex conformational rearrangements are considered as in the system presented here. Neural networks, on the other hand, are particularly suited for this task. For example, in the pioneering work by Ma and Dinner [27] neural networks were used to predict the committor based on a set of coordinates for the isomerization of alanine dipeptide. More recently, Jung et al. proposed and implemented a combination of path sampling and deep neural networks to guide the sampling and to extract the reaction coordinate [25, 26, 52]. Following the work by Jung et al. [25, 26], we used a deep neural network to learn the reaction coordinate from the outcome of the committor analysis by minimizing the likelihood loss function [53, 54]. Note however, that we did not guide the sampling as described in ref. [25, 26]. Instead, we performed a preceding committor analysis (as described above) and use the committor to map the molecular conformations onto the reaction coordinate *q*^pred^(**X**). Each conformation was described by a set of 83 physical properties (referred to as features **X** in the following). The committor was parametrized as in ref. [25, 26]

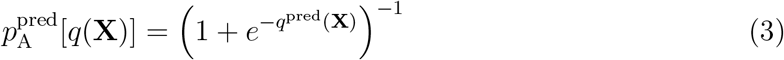

where the predicted committor *p*^pred^ is a nonlinear function of all the features **X** of the system. Note that by rescaling the reaction coordinate via 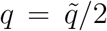 the frequently used expression for the committor by Peters and Trout [54] is retrieved:

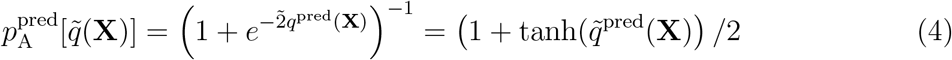

Therefore, eq. 3 and 4 are expected to yield identical results. The likelihood that a model can reproduce the observed data is given by [25, 26, 53, 54]

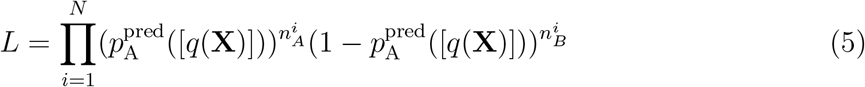

where *N* is the number of shooting points and 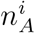 and 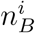 are the number of trajectories that reach state A and B from shooting point *i*, respectively. Following the work by Jung [25, 26], we modeled the committor with a deep neural network. Hereby, the weight matrix that defines the connection between the nodes of the deep neural network was optimized by minimizing the negative log likelihood loss *l* [25, 26]

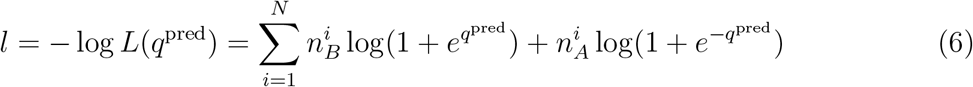

The overall accuracy and generalization of the machine learning model (i.e. the deviation of the predicted values 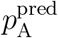 from the simulated ones 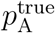) was estimated by dividing the available data into a training set, a validation set and a test set. The first was used to train the model, the second to validate the training result. The third was used to check whether the final model is able to correctly predict the values for structures that were not used for training.

### Training a deep learning regression model with hyperparameter optimization

One challenge in the application of neural networks is to choose the model architecture. As illustrated in Figure 1A, the accuracy of the network to predict the commitment probability of a conformation and to autonomously select the transition state strongly depends on the underlying model architecture. However, there is no generic way to determine all the model parameters such as the number of hidden layers, the number of neurons and all hyperparameter a priori. Moreover, a network architecture that works well for one system might fail to capture other systems or dissimilar problems. Therefore, building an optimal deep neural network manually by trial and error is a time consuming problem which highly depends on human expertise and intuition. In order to make this progress more efficient, we developed an algorithm that automatically finds the optimal network architecture for any given problem. First, we specified a class of multilayer perceptron (MLP) regression models that are described by a set of hyperparameter [55]. Subsequently, we selected the optimal model by running a Keras Tuner random search hyperparameter optimization [56]. Hereby, the generic MLP was defined by the following set of hyperparameter: number of hidden dense layers (1-6), number of neurons per hidden layer (32-256), activation function (Rectified Linear Units (ReLU) or Scaled Exponential Linear Unit (SELU)), position of a single dropout layer (none/after feature input/middle layer/before output layer), dropout rate (0-40%), and learning rate for the Adam Optimizer (logarithmic sampling from 10^−4^ - 10^−2^). For ReLU activation functions, a Glorot uniform initialization and a normal dropout layer were used. For SELU activation functions, a Lecun normal initialization and an alpha dropout layer were used. ReLU and SELU are the standard choices for regression models in Keras and were therefore used. The custom loss function by Jung et al. [25, 26] according to Eq. 6 was used. All models were trained via back propagation using the stochastic gradient algorithm.

**FIG. 1:**
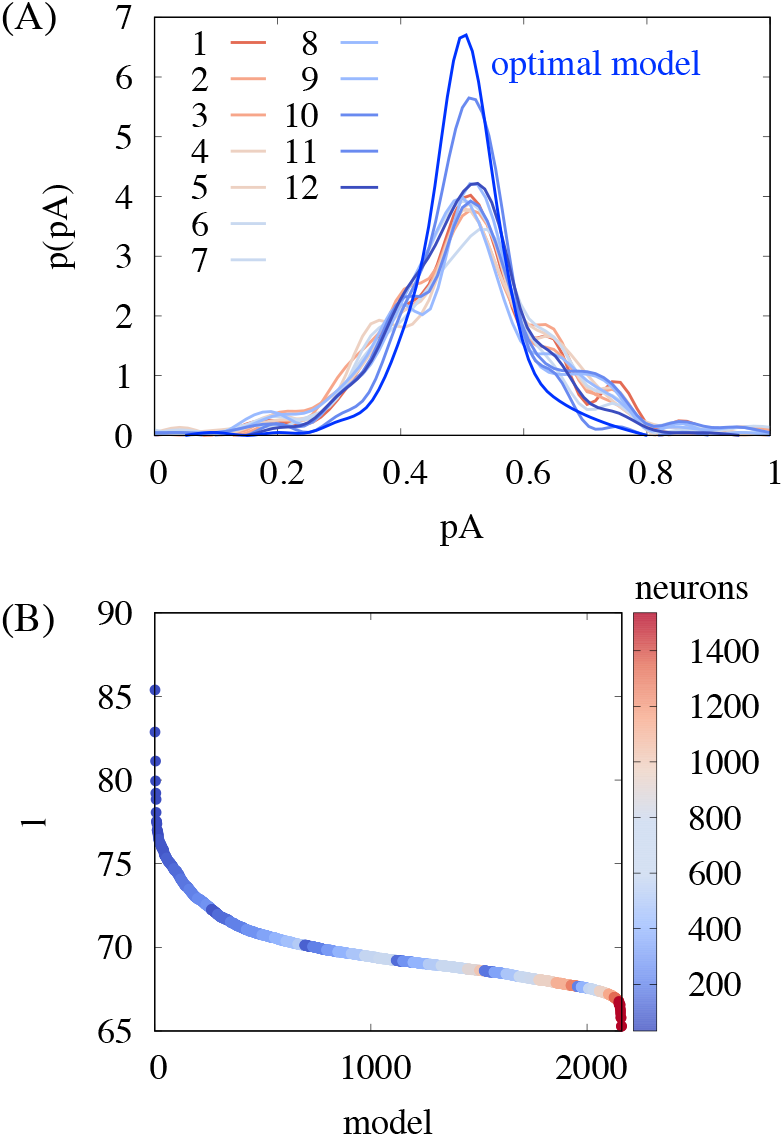
Dependence of the performance on the model architecture. (A) Comparison of the committor distribution 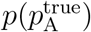 for transition states selected from deep learning algorithms with different model architectures (see Supporting Information for further details of the models). (B) Validation likelihood loss *l* for a random subset of model architectures. One parameter of the architecture, the total number of neurons, is shown in color. In total, several thousand different model architectures were explored by running a Keras Tuner random search in the space of model parameters.

The data was split into 20% test set, 72% training set and 8% validation set. All features were normalized using the Keras standard scaler with mean *μ* = 0 and standard deviation *σ* = 1. Each model was set up for 50 epochs with a training batch size of 128. Early stopping with patience 10 was used on the validation loss. Multiple thousand different model architectures were explored and the likelihood loss of a random subset of 2,160 models is shown in Figure 1B.

For further optimization, a subset of best models based on minimal validation loss was chosen. The training of the best model subset was continued for up to 200 epochs with the previous batch size and early stopping. In addition, a stepwise reduction of the learning rate by a factor of 5 down to 10^−5^ was used if the validation loss reached a plateau for 7 training steps. Finally, the best model was chosen based on the highest maximum in the 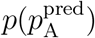 distribution.

In summary, the optimization algorithm allows us to find the optimal model architecture in a quick and reliable fashion. The optimal deep learning model has 5 hidden layers with 256 neurons, ReLU activation and an initial learning rate of 1.9 ∗ 10^−4^. A dropout layer with 30% dropout is placed directly before the output layer. All code was developed with Keras [57], scikit-learn [58] and tensorflow [59].

### Features

For each shooting point, we calculated 83 features that reflect different structural properties of the molecular conformation. Focusing only on the dominant indirect exchange, the dataset has 17,477 entries. Each entry consists of 83 features and the label 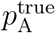 calculated for each shooting point using the committor analysis. Specifically, the features include all distances *r*_*i*_ between Mg^2+^ and the 20 closest water molecules, distances between Mg^2+^ and different RNA atoms and the Cl^−^ ion, all angles *a*_*i*_ formed between Mg^2+^, O1P and the 20 closest water molecules, the Steinhardt-Nelson order parameters *q*_3_, *q*_4_, *q*_6_ [60] of the first and second hydration shell, tetrahedral order parameters [46], and the number of hydrogen bonds in the first and second hydration shell and between RNA and water (Table S1).

In addition, a putative reaction coordinate *λ* based on human expert knowledge was defined and includes the Mg^2+^-RNA distance *r*_*I*_ and the effective hydration distance *s*_6_ via a trigonometric function

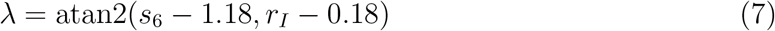

### Feature relevance

To this end, the machine learning model employs a large number of features to model the reaction coordinate and to predict the commitment probability 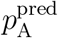 of new structures. In the following, we aim to determine the importance of each input feature. Since the deep learning network consists of a large number of nodes, weighted sums and nonlinear transformations, the feature relevance was calculated numerically by selectively replacing single features or combinations of features by noise and resampling the model.

For the single feature relevance, we generated for every input feature *i* a dataset in which the values **X**_*i*_ were replaced by randomly permuted values 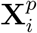. The random permutation proach was chosen as it exactly conserves the original distribution. The normalized relevancaep-of the *i*-th feature 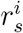 was defined as

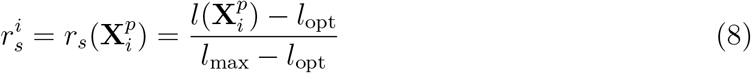

where *l*_opt_ is the converged loss of the optimal model and *l*_*max*_ is the largest value upon permutation. The relevance corresponds to the loss of information (i.e. the negative log likelihood increase) upon effectively removing a single feature from the dataset while leaving all other features unchanged. Accordingly, the single feature rank ranges from low (unimportant) to high (important).

For single feature permutations importance, standard libraries can be used [58, 61]. For this work, we implemented a custom algorithm that extends the standard permutation importance algorithms to be able to measure the importance of combined feature sets (see below). Note that an alternative approach for the single feature analysis was proposed in ref. [25] where uniform random noise was used instead of random permutations.

For the relevance of combined features, *N* (*N* + 1)*/*2 = 3486 permuted datasets were generated as follows. Starting from the optimal model, the feature which yielded the smallest loss upon permutation was removed (rank 1). This feature contained the smallest amount of information and was consequently least important for the reaction. Maintaining the permutation of rank 1, we selected the second feature that yielded the smallest loss upon permutation (rank 2). The procedure was repeated until all features are permuted. The normalized relevance for effectively removing the information of *n* features is given by

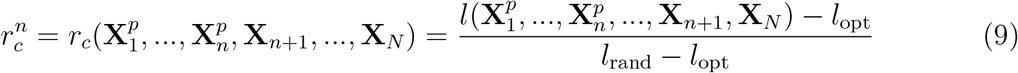

where *l*_opt_ is the converged loss of the optimal model and *l*_rand_ is loss obtained by permuting all features. Here, the relevance corresponds to the increase of the negative log likelihood upon removing the combination of *n* features that contain the least amount of information. Accordingly, the feature combination rank ranges from low (least important combinations) to high (most important combination).

## RESULTS AND DISCUSSION

### Free energy landscape of Mg^2+^-RNA interactions

During association, one of Magnesium’s six strongly bound hydration waters is removed to facilitate a direct contact between Mg^2+^ and the phosphate oxygen. In the simplest case, this ligand exchange can be described by the distance *r*_*I*_ between Mg^2+^ and the phosphate oxygen while all other degrees of freedom are integrated out. The corresponding one-dimensional free energy profile is shown in Figure 2A. The free energy profile has two stable states. State A corresponds to the inner-sphere conformation, state B correspond to the outer-sphere conformation. The free energy barrier from outer-sphere to inner-sphere is about 21 *k*_B_*T* and exactly matches the value for water exchange [30]. Consequently, the barrier corresponds to the free energy necessary to remove one water molecule from the first hydration shell in order to facilitate a direct contact with the phosphate oxygen. The inner-sphere conformation is thermodynamically more stable in agreement with experimental findings [18] and Collins’ empirical rule like-seeks-like [62]: Due to the high charge density of the phosphate oxygens, direct ion pairing with a high binding affinity is expected.

**FIG. 2:**
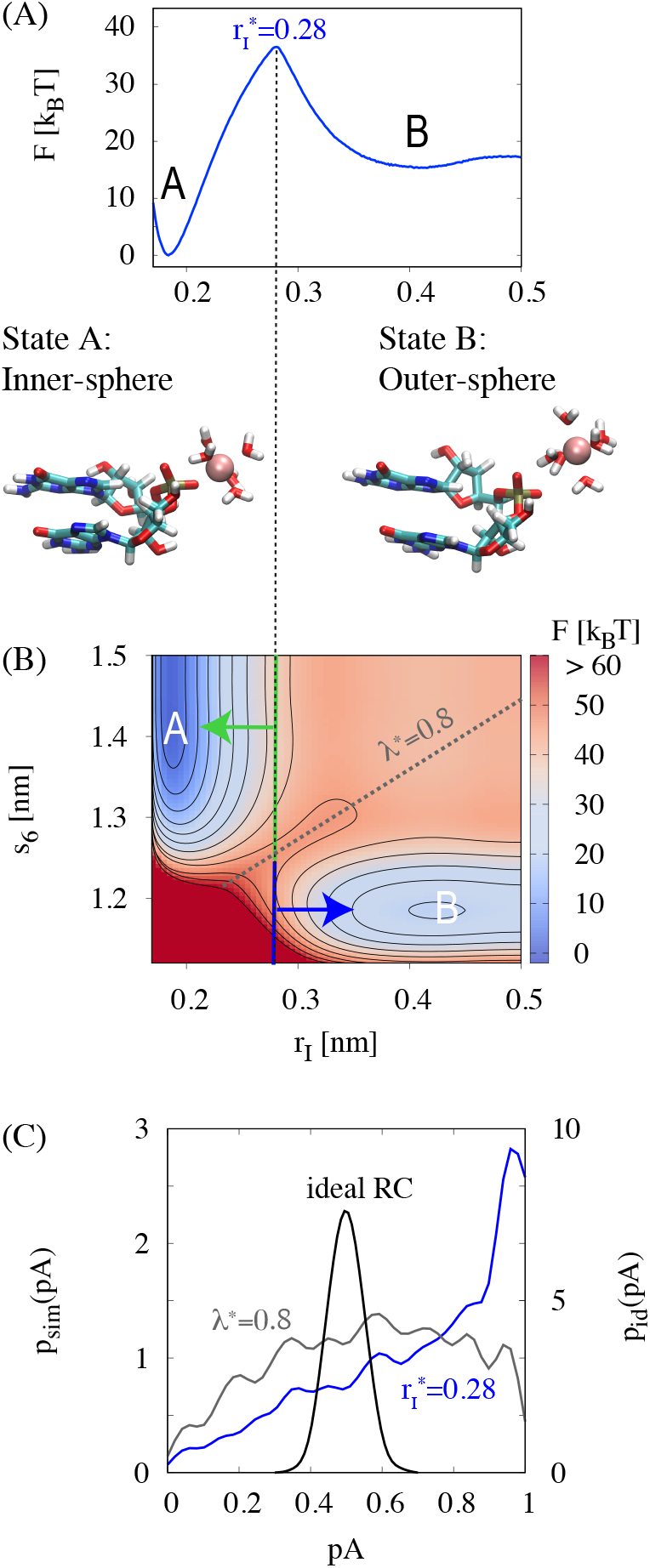
Free energy landscape of Mg^2+^-RNA interactions. (A) Free energy profile *F* (*r*_*I*_) as function of the distance between Mg^2+^ and the phosphate oxygen. Simulation snapshots of the two stable states are shown. (B) Two-dimensional free energy landscape *F* (*r*_*I*_, *s*_6_) as a function of *r*_*I*_ and *s*_6_ where *s*_6_ is the sum over the distances between Mg^2+^ and the 5 closest and the exchanging water molecule. The dashed line indicates conformations from the top of the free energy profile with 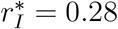 shown in (A). Trajectories, initiated from the upper green stripe, relax back into state A, trajectories from the lower blue stripe relax back into state B. The diagonal dashed line for *λ*\* = 0.8 approximates the saddle between states A and B. (C) Committor distributions *p*_Sim_(*p*_A_) for trajectories initiated with 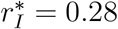 (top of the one-dimensional free energy profile shown in A) and with *λ*\* = 0.8 (saddle of two-dimensional free energy profile shown in B). *p*_id_(*p*_A_) is the distribution expected for an ideal reaction coordinate with 3500 shooting points.

Using transition state theory TST (Eq. 2), we can provide an upper estimate of the exchange rate or conversely a lower estimate of the lifetime. Here it is essential to mention that the true rate could be significantly smaller due to the violation of the non-recrossing hypothesis. As lower limit, Mg^2+^ is estimated to remain about 300 s in the inner-sphere conformation before transitioning to outer-sphere. Similarly, Mg^2+^ is estimated to remain about 0.2 ms in the outer-sphere conformation before transitioning back to inner-sphere. Note that the exchange between the outer-sphere conformation and bulk is much faster since the free energy barrier is significantly smaller (Figure 2A). The lifetime of the outer-sphere conformation is in agreement with the millisecond timescale observed experimentally [19, 21, 22]. On the other hand, the computed lifetime of the inner-sphere conformation is likely too high reflecting the shortcoming of current Mg^2+^ force fields in reproducing experimental binding affinities at nucleic acid binding sites [24, 63, 64].

The quality of a reaction coordinate can be assessed by a committor analysis. For an ideal reaction coordinate, about half of the trajectories initiated from the barrier top are expected to relax back to either stable state. Consequently, the distribution *p*(*p*_A_) of the probability to relax back to state A should have a sharp peak at *p*_A_ ≈ 1/2 (binomial committor distribution). While *r*_*I*_ provides a simplified description of the reaction, it is not an adequate reaction coordinate by itself. The committor analysis (Figure 2C) shows that most conformations, initiated with *r*_*I*_ = 0.29 relax back to the inner-sphere conformation. Therefore, the Mg^2+^-oxygen distance alone is insufficient to describe the dynamics of the transition. To provide a more realistic picture, the water molecules from the first hydration shell have to be included. Figure 2B shows the two-dimensional free energy landscape as a function of *r*_*I*_ and the hydration parameter *s*_6_, which includes the distances of the five closest waters and the exchanging water (Eq. 1). From the two-dimensional representation, the failure of *r*_*I*_ as reaction coordinate can be rationalized: Trajectories starting from the upper panel (*s*_6_ > 1.24 nm) are committed to state A, while trajectories starting from the lower panel (*s*_6_ < 1.24 nm) are committed to state B. *r*_*I*_ and *s*_6_ can be combined into a putative, human knowledge-based reaction coordinate *λ* (Eq. 7). Based on the committor distribution (Figure 2C), *λ* provides significant improvement compared to *r*_*I*_. Still, conformations with *λ*\* = 0.8 lead to a much broad distribution compared to the ideal binomial distribution.

These results show, that free energy profiles along a few simple coordinates provide valuable initial insight. Yet, the committor analysis reveals that the exchange dynamics is more complex than the free energy profiles might suggest.

### Kinetic pathways from transition path sampling

To gain insight into the kinetic pathways of Mg^2+^ association and dissociation, transition path sampling is applied to sample a large number of inner-to-outer sphere transitions. Four representative transition paths that connect the two stable states are shown in Figure 3A. Figure 3C shows the distribution of transition times. Typically, the exchanging water molecules spend less than 0.5 ps in transition. This time is considerably smaller compared to the millisecond timescale of the stable states. The clear separation of timescales highlights the path sampling method as a particularly powerful sampling strategy for these systems.

**FIG. 3:**
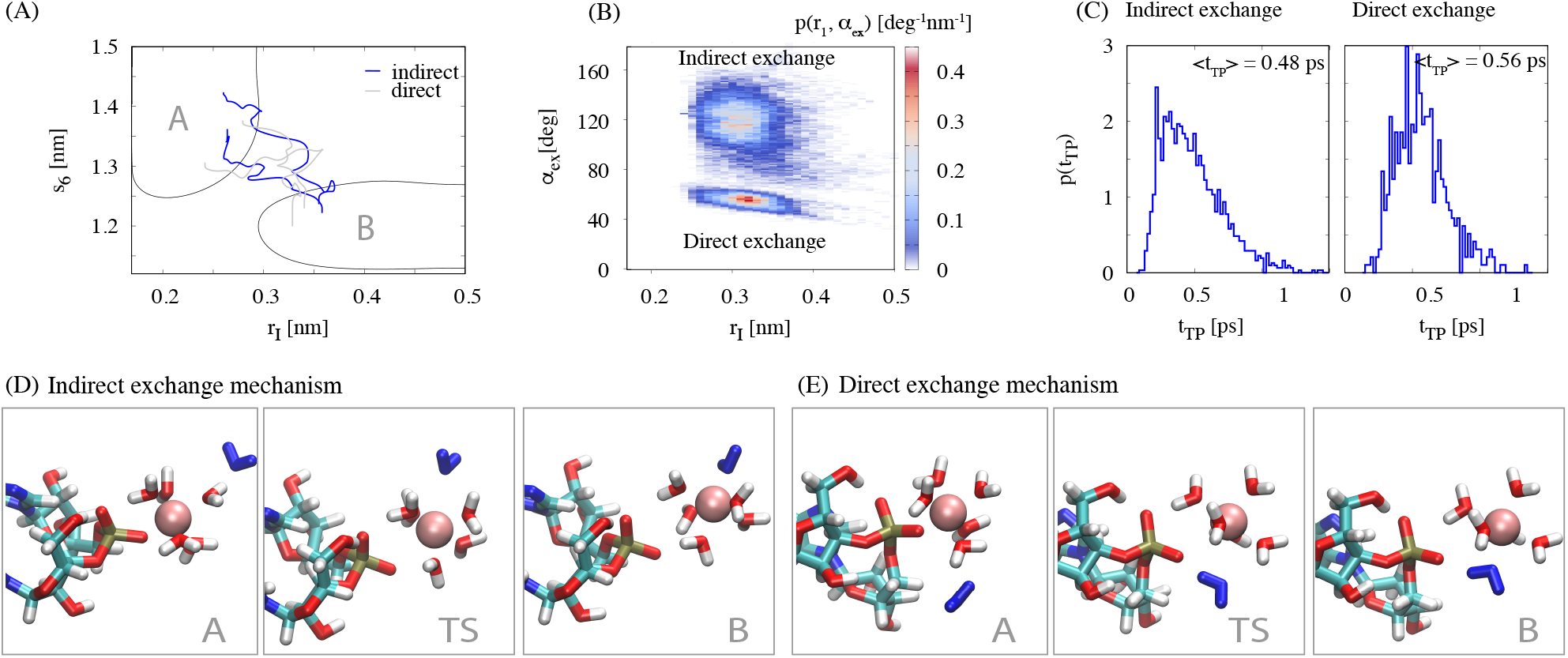
Kinetic pathways from transition path sampling. (A) Four representative pathways connecting the two stable states. Blue/gray pathways correspond to the indirect/direct exchange mechanism. (B) Probability distribution along transition pathways for the distance r_*I*_ between Mg^2+^ and the phosphate oxygen and the exchange angle *α*_ex_ between phosphate oxygen, Mg^2+^ and exchanging water oxygen. (C) Distribution of transition times for the indirect and the direct exchange mechanism. (D) Indirect Mg^2+^ exchange mechanism in which the leaving phosphate oxygen and the incoming water molecule occupy different positions on the water octahedron. (F) Direct Mg^2+^ exchange mechanism in which the leaving phosphate oxygen and the incoming water molecule occupy the same position on the water octahedron. In (D) and (E) the five closest water molecules are shown. The incoming water molecule is highlighted in blue.

The distribution of the exchange angles (Figure 3B) along reactive pathways indicates that two alternative exchange pathways exist: In the *indirect exchange mechanism*, the leaving oxygen ligand and the entering water molecule occupy different positions on the water octahedron (Figure 3D). During activation, one water molecule from the second hydration shell enters the molecular void between the hydration shells. This motion leads to concerted motion of the phosphate oxygen out of the first hydration shell. The distances of the leaving phosphate oxygen and the entering water are elongated compared to their equilibrium values (Table I). Reflecting balanced electrostatic interactions, the Mg^2+^-phosphate oxygen distance is smaller than the Mg^2+^-water oxygen distance due to the smaller partial charge on the phosphate oxygen compared to the water oxygen. The distances of the five closest molecules remain relatively unchanged (Table I). Yet, they rearrange such that the transition state has an approximate mirror symmetry. The mirror plane is perpendicular to the plane composed of Mg^2+^, the phosphate and water oxygen and contains three water molecules (Figure 3D).

**TABLE I:**
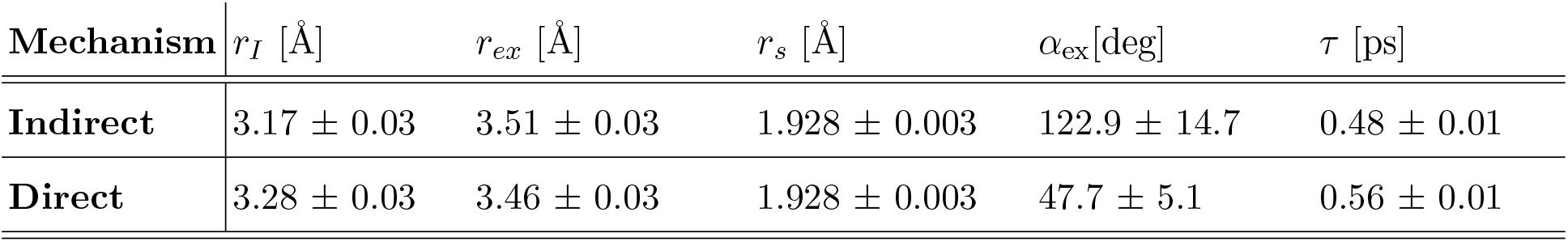
Properties of the transition state ensemble: Mg^2+^-O1P distance *r*_*I*_, Mg^2+^-oxygen distance of exchanging water *r*_*ex*_, Mg^2+^-oxygen distance of the five closest water molecules *r*_*s*_, angle between O1P, Mg^2+^ and the exchanging water molecule *α*_ex_, average transition time *τ*. Standard deviations are indicated. A sample size of 1759 or 754 transition states was used for the indirect and direct exchange mechanism, respectively.

| **Mechanism** | $r_I$ [Å] | $r_{ex}$ [Å] | $r_s$ [Å] | $\alpha_{ex}$[deg] | $\tau$ [ps] |
| --- | --- | --- | --- | --- | --- |
| **Indirect** | $3.17 \pm 0.03$ | $3.51 \pm 0.03$ | $1.928 \pm 0.003$ | $122.9 \pm 14.7$ | $0.48 \pm 0.01$ |
| **Direct** | $3.28 \pm 0.03$ | $3.46 \pm 0.03$ | $1.928 \pm 0.003$ | $47.7 \pm 5.1$ | $0.56 \pm 0.01$ |

In the *direct exchange mechanism*, the leaving oxygen ligand and the entering water molecule occupy the same positions on the water octahedron (Figure 3E). The exchange arises via the attack of the incoming water onto the edge of the octahedron formed by the oxygen ligands. Similar as in the indirect pathway, the Mg^2+^-phosphate oxygen distance is slightly smaller than the Mg^2+^-water oxygen distance while the distances of the five closest water molecules remain relatively unchanged (Table I). In the transition state, the five closest water molecules form a square pyramidal coordination and the transition state has a distorted C_2_ symmetry (Figure 3E).

Based on the change of bond length during activation, both pathways correspond to an interchange dissociative (I_*d*_) process and are akin to the pathways of water exchange [30]. In equilibrium, the indirect exchange is observed much more frequently (92%) compared to the direct mechanism (8%) since conformations with cis positions of exchanging ligands (direct pathways) are energetically less favorable compared to trans positions (indirect pathway) [65].

The results reveal that phosphate oxygen, exchanging water and the five closest water molecules play a decisive role in the exchange mechanism. However, additional simulations in which those coordinates were fixed while the solvent outside the first hydration shell was relaxed, show that they are yet insufficient to predict *p*_*A*_. Consequently, solvent molecules beyond the first hydration shell are crucial for the exchange reaction and need to be considered explicitly.

### Optimized deep neural networks for quantitative predictions

During the transition from outer-to-inner sphere binding, water molecules from the first two hydration shells couple to the exchange. The reordering of close and distant water molecules determines whether the reaction can proceed or not. This behavior might be expected since the kosmotropic Mg^2+^ ion causes strong orientational ordering in the first hydration shells leading to long-ranged and collective interactions. However, resolving the subtle rearrangements and providing a quantitative description of the solute-solvent coupling is a demanding problem that is impossible to solve by visual inspection. In order to make progress in quantifying the solvent participation, we use a deep neural network to model the outer-to-inner sphere exchange reaction. Here, we focus the machine learning on the dominant indirect exchange. In particular, we use 17,477 structures along independent transition paths and their commitment probabilities *p*_A_ (Figure 4A) to learn the functional relation between *p*_A_ and the features describing the structure of Mg^2+^, RNA and the first two hydration shells.

**FIG. 4:**
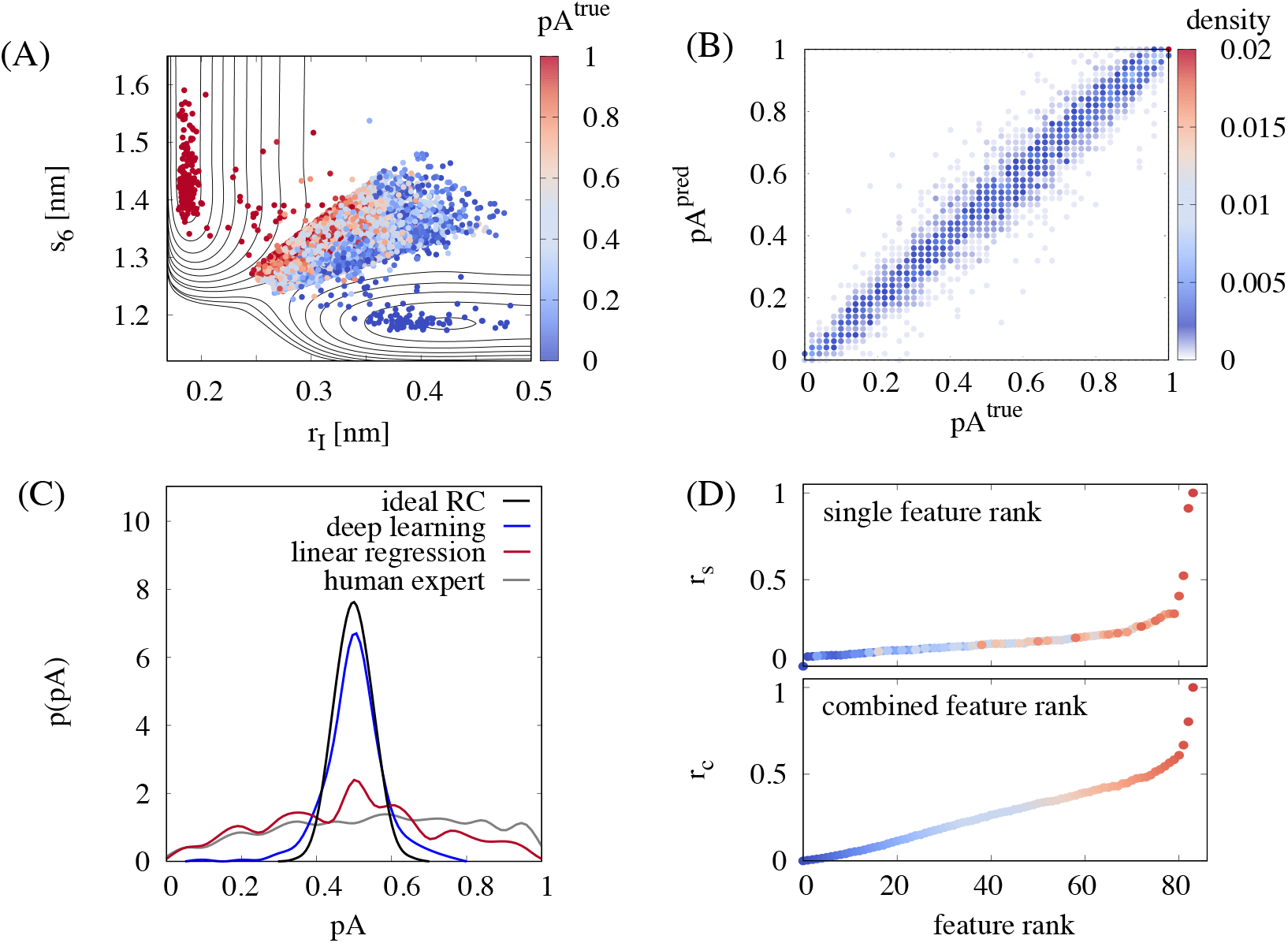
Reaction coordinate from artificial intelligence. (A) Dataset used for machine learning. The committor probability 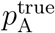 and free energy contour is shown as function of the features *r*_*I*_ and *s*_6_ for the full dataset consisting of 17,477 entries for indirect exchange pathways (72% training set, 8% validation set, 20 % test set). (B) Committor values 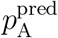 predicted by the optimized deep neural network correlated with the values 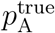 obtained from the committor simulations. The RMS error of the prediction is 0.065. (C) Comparison of the committor distribution 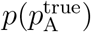 for transition states selected from the deep learning algorithm, from linear regression over all features, from the human expert knowledge using Eq. 7 and the binomial distribution expected for an ideal reaction coordinate. (D) Ranking of the features according to their relevance in the machine learned reaction coordinate: Single feature relevance *r*_*s*_ (top) and combined feature relevance *r*_*c*_ (bottom). The color indicates the rank of the features according to the combined feature relevance. The corresponding features are shown in Table II and Table S2.

The choice of the architecture of the deep learning model is essential for its performance. To derive the optimal deep learning model, we have defined an algorithm that systematically selects the optimal architecture by scanning through thousands of individual models using hyperparameter optimization. In order to make robust predictions and prevent overfitting, we optimize the neural network on the training set and select the model that performs best on the validation set. Finally, using the test set, we illustrate the performance of the optimal machine learning model in predicting the commitment probability of unknown structures (Figure 4B). The results demonstrate that the optimized deep neural network is capable of predicting *p*_A_ with an RMS error of 0.065. The high accuracy in predicting the process of the reaction clearly shows that the network extracts all relevant information, combines it in a weighted nonlinear fashion to a scalar reaction coordinate and captures the details of the solute-solvent coupling precisely.

Further insight into the quality of the predictions is obtained from the distribution of commitment probability *p*(*p*_A_) of the transition states selected autonomously by the neural network (Figure 4C). The distribution is unimodal and sharply peaked at *p*_A_ ≈ 0.5 and closely resembles the binomial distribution expected for an ideal reaction coordinate. Consequently, the information contained in the features is sufficient for the network to single out the transition state. The results from deep learning provide significant improvement compared to the reaction coordinate based on human knowledge and highlight the machine learning ansatz as a particularly useful strategy in recognizing the relevant patterns. In addition, the results from deep learning outperform the results from linear regression based on the exact same features (Figure 4C). Therefore, hidden layers and a nonlinear combination of features using activation functions are essential to describe the many-body interactions. At the same time, the complexity of the deep learning model prohibits further insight into the reaction mechanism. In this regard, it is particularly useful to evaluate the contribution of each feature. Two complementary rankings are presented in Figure 4D. The single feature ranking quantifies the relevance of a single feature independent of all others. Since the single feature ranking can contain redundant information, the complementary combined feature rank is given, which yields the most relevant combination of features. Taken together, a consistent picture emerges: All features describing the molecular structure within the first two hydration shells contain information that is relevant for the reaction. Adding this information to the model leads to a continuous decrease of the likelihood loss and therefore to a linear increase of the relevance *r*_*c*_. Finally, four features give rise to an exponential increase in the relevance and carry about 40% of the information (Table II). These features reflect the concerted motion of the leaving phosphate oxygen and the entering water molecule as well as the rearrangement of the 5 closest water molecules from octahedral to mirror symmetry. At the same time, the remaining 60% of the information is distributed over the remaining 79 features reflecting the importance and many-body nature of the solvent-solute coupling.

**TABLE II:** Ranking of the 8 most important features in the machine learned reaction coordinate according to their normalized relevance (Eq. 8, 9). The combined feature rank (c-rank) corresponds to the combination of *N* − *n* features that contains most information i.e. yields the smallest log likelihood loss upon permuting *n* features. The single feature rank (s-rank) corresponds to the feature which yields the highest loss upon permuting the feature while leaving all other features unchanged. The ranking of all *N* = 83 features is given in Table S1.

| **c-rank** | feature | **s-rank** | feature |
| --- | --- | --- | --- |
| 84 | $r_I - r_{\text{ex}}$ | 84 | $r_I - r_{\text{ex}}$ |
| 83 | $r_I$ | 83 | $r_I$ |
| 82 | $q_{40}$ | 82 | $q_{40}$ |
| 81 | $r_{\text{P}}$ | 81 | $r_{\text{P}}$ |
| 80 | $h_{\text{O1P}}^1$ | 80 | $h_{\text{ex}}^1$ |
| 79 | $h_{\text{ex}}^1$ | 79 | $q_{41}$ |
| 78 | $a_1$ | 78 | $r_{\text{Cl}}$ |
| 77 | $s_6$ | 77 | $a_7$ |

## CONCLUSION

Characterizing the kinetic pathways of Mg^2+^ binding to biomolecules such as RNA is fundamental in understanding its functional role in biochemical processes. Yet, the transition from water-mediated outer-sphere to direct inner-sphere binding is on the millisecond timescale and therefore out of reach for conventional all-atom simulations. To fill this gap, we used transition path sampling to resolve the kinetic pathways of Mg^2+^ binding to the most important ion binding site on RNA, namely the phosphate oxygen. The results reveal a superior indirect and an inferior direct exchange pathway. In both pathways, the molecular void in the first hydration shell provoked by the leaving phosphate oxygen is immediately filled by an entering water. At the same time, the water molecules from the first and second hydration shell couple to the concerted exchange. These long-ranged and collective interactions are a direct consequence of the high ionic charge density of Mg^2+^ which provokes strong ordering in the surrounding solvent and an intimate coupling between solute and solvent. Consequently, ligand exchange gives rise to a complex interplay of orientational, packing and hydrogen bonding effects in which the collective reorientation and translation of several solvent molecules becomes important.

A complete understanding of the reaction dynamics requires knowledge of the detailed molecular motions involved in the exchange process. However, quantifying their contribution is exceptionally difficult. Instantaneous fluctuations, resulting from different solvent conformations, influence the progress of the reaction in particular since the timescale of solvent reorientation (about 10 ps) is slower than the reactive motion itself (0.4 ps). In order to quantify how the reactant and solvent dynamics are coupled, the rapidly growing field of machine learning offers exciting possibilities in extracting complex patterns in large datasets and in finding optimal reaction coordinates [25–27, 52, 66, 67]. The results presented here reveal that a properly optimized deep neural network is particularly suited to recognize the molecular motions that occur during the course of the reaction, to extract the relevant information, and to predict the commitment probability with high accuracy. About half of the information on the reaction dynamics is contained in the concerted motion of the leaving phosphate oxygen and the entering water molecule as well as in the rearrangement of the 5 closest water molecules. The other half is contained in the solvent structure rendering deep neural networks particularly useful in recognizing the relevant molecular structures.

The question how the solvent affects reactions in aqueous solutions is ubiquitous in all areas of chemistry and biology. Here, machine learning provides a promising perspective to explore the intimate coupling between charged solutes and the solvent at the molecular level [25, 26, 66].

## SUPPORTING INFORMATION

Further discussion of the two-dimensional free energy landscape, the commitment probability from constrained simulations, feature definition and the feature ranking.

## ACKNOWLEDGMENTS

This work was funded by the Deutsche Forschungsgemeinschaft (DFG, German Research Foundation), Emmy Noether program, Grant No. 315221747. LOEWE CSC and GOETHE HLR are acknowledged for supercomputing access. N.S. thanks Roland K. O. Sigel and Harald Schwalbe for inspiring discussions on metal ion binding. N.S. also thanks Phill L. Geissler and Marieke Schor for helpful discussion on transition path sampling. In addition, N.S. thanks Hendrik Jung and Roberto Covino for fruitful discussions and for comments on the manuscript. Most importantly, we thank Gerhard Hummer for introducing us to the combination of path sampling and machine learning, for critical comments on the manuscript and stimulating discussions.

## SUPPORTING INFORMATION

### 1 Free energy landscape

The two-dimensional free energy landscape as a function of the distance between Mg^2+^ and the phosphate oxygen O1P is calculated from umbrella sampling. Without further restraints, the system has three stable states (Figure S1). State A corresponds to the inner-sphere conformation in which Mg^2+^ is coordinated by the O1P atom and five additional water molecules. State B corresponds to the outer-sphere conformation in which the O1P atom is replaced by the selected water molecule *W*_ex_ for which the biasing umbrella potential is applied. State C corresponds to conformations in which O1P is replaced by any other of the *N* − 6 water molecules (where *N* is the number of water molecules in the simulations box and 6 is the coordination number). In this two-dimensional projection, transitions between state A and state C involve additional water molecules that are hidden in the selected two-dimensional projection. Here, an *N* − 6 dimensional representation would be required to capture all relevant coordinates. However, since all outer-sphere states are identical since the water molecules are indistinguishable, it is sufficient to show only the exchanges between state A and state B in the two-dimensional projection. This is achieved by applying an additional biasing potential on the coordination number *n*_1_. Hereby, the coordination number is calculated from the switching function

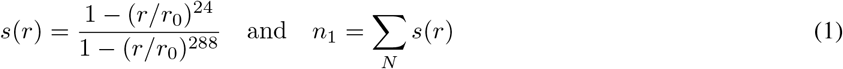

with the ion-water distances *r* and the cutoff *r*_0_ = 0.3 nm. With the hydration bias, all water molecules other than the five closest water molecules and the selected exchanging water are forced to remain outside the first hydration shell by setting *n*_1_ = 0 using a force constant of *k*_*h*_ = 1000 kJ/mol.

**Figure S1:**
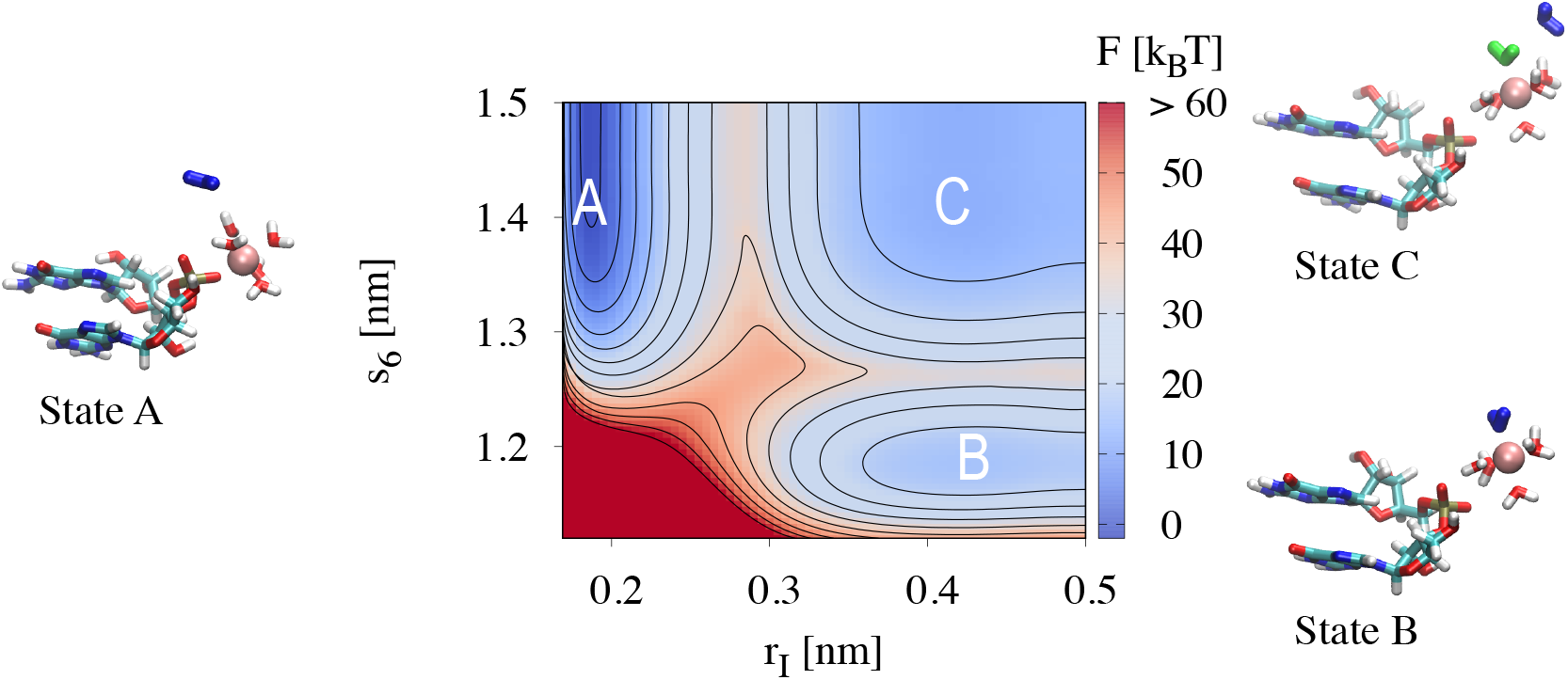
Two-dimensional free energy landscape *F*(*r*_*I*_, *s*_6_) of the Mg^2+^-O1P distance *r*_*I*_ and the hydration parameter *s*_6_. *s*_6_ is the sum of the distances to the five closest water molecules (shown in red and white) and the selected water molecule *W*_*ex*_ for which the biasing umbrella potential is applied (shown in blue). The green water molecule indicates one of the *N* − 6 unlabeled water molecules. The energy contour spacing is 5 *k*_B_T.

### 2 Dependence of the performance on the deep learning model architecture

The quality of the deep learning model strongly depends on the choice of the underlying model architecture. To illustrate the dependence of the accuracy on the deep learning model architecture, 12 architectures out of several thousands were used to predict the transition state 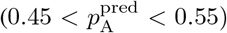 and to calculate the resulting committor distribution 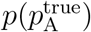. The model architectures selected for this analysis were the following:

<u>Optimal model:</u> 5 hidden layers with 256 neurons, ReLU activation and an initial learning rate of 1.9 ∗ 10^−4^. A dropout layer with 30% dropout is placed directly before the output layer.
<u>Model 1:</u> 5 hidden layers with 256 neurons, ReLU activation and an initial learning rate of 2.8 ∗ 10^−3^. A dropout layer with 30% dropout is placed after the second hidden dense layer.
<u>Model 2:</u> 3 hidden layers with 192 neurons, ReLU activation and an initial learning rate of 1.6 ∗ 10^−3^. A dropout layer with 20% dropout is placed after the input layer.
<u>Model 3:</u> 4 hidden layers with 192 neurons, ReLU activation and an initial learning rate of 3.8 ∗ 10^−3^. A dropout layer with 20% dropout is placed after the input layer.
<u>Model 4:</u> 3 hidden layers with 192 neurons, ReLU activation and an initial learning rate of 5.9 ∗ 10^−3^. A dropout layer with 10% dropout is placed after the input layer.
<u>Model 5:</u> 4 hidden layers with 224 neurons, ReLU activation and an initial learning rate of 1.3 ∗ 10^−4^. A dropout layer with 30% dropout is placed after the input layer.
<u>Model 6:</u> 5 hidden layers with 256 neurons, ReLU activation and an initial learning rate of 1.3 ∗ 10^−3^ without dropout layer.
<u>Model 7:</u> 5 hidden layers with 192 neurons, ReLU activation and an initial learning rate of 8.0 ∗ 10^−4^. A dropout layer with 20% dropout is placed after the input layer.
<u>Model 8:</u> 4 hidden layers with 224 neurons, ReLU activation and an initial learning rate of 4.3 ∗ 10^−4^. A dropout layer with 20% dropout is placed after the second hidden dense layer.
<u>Model 9:</u> 5 hidden layers with 256 neurons, SELU activation and an initial learning rate of 3.6 ∗ 10^−3^ without dropout layer.
<u>Model 10:</u> 6 hidden layers with 256 neurons, ReLU activation and an initial learning rate of 4.0 ∗ 10^−3^ without dropout layer.
<u>Model 11:</u> 3 hidden layers with 224 neurons, ReLU activation and an initial learning rate of 2.0 ∗ 10^−3^ without dropout layer.
<u>Model 12:</u> 2 hidden layers with 192 neurons, ReLU activation and an initial learning rate of 5.3 ∗ 10^−3^. A dropout layer with 30% dropout is placed after the first hidden dense layer.

### 3 Commitment probability from constrained simulations

To test the influence of the second hydration shell on the commitment probability, we performed additional constrained simulations. For 600 shooting points, the positions of Mg^2+^, RNA, five closest water molecules and exchanging water were kept constant. All other atoms were equilibrated for 1 ns in the NVT ensemble. Subsequently, 100 trajectories were initiated with velocities drawn from a Maxwell-Boltzmann velocity distribution and run forward and backward for 2 ps (see main manuscript for further details). The commitment probability *p*_A_ was calculated from the fraction of trajectories that reached state A. Figure S2 shows a comparison of the commitment probability for 600 conformations drawn directly from transition path sampling and for the same conformations after the constrained equilibration described above. Clearly, the values of *p*_A_ are different for the unconstrained and the constrained conformations. We conclude that the positions of Mg^2+^, RNA, five closest water molecules and exchanging water are insufficient to predict *p*_A_. Consequently, the exchange reaction is influenced by water molecules outside the first hydration shell. A quantitative description should therefore include the water molecules from the second hydration shell or even beyond.

### 4 Features

Table S1 lists all features used for deep learning. The features include all distances *r*_*i*_ between Mg^2+^ and the 20 closest water molecules, distances between Mg^2+^ and different RNA atoms and the Cl^−^ ion, all angles *a*_*i*_ formed between O1P, Mg^2+^ and the 20 closest water molecules, Steinhardt-Nelson order parameters, tetrahedral order parameters and numbers of hydrogen bonds for different groups of atoms. Hereby, the tetrahedral order parameter calculates the degree to which the water molecules around O1P or W_*ex*_ have a tetrahedral order. It is defined as

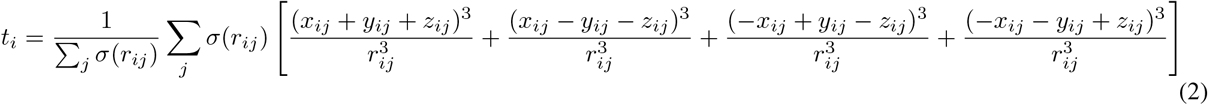

where *r*_*ij*_ is the magnitude of the vector connecting atom *i* and atom *j* and *x*_*ij*_, *y*_*ij*_ and *z*_*ij*_ are its three components. *σ*(*r*_*ij*_) is a switching function with a cutoff of *r*_0_ = 0.35 nm.

**Figure S2:**
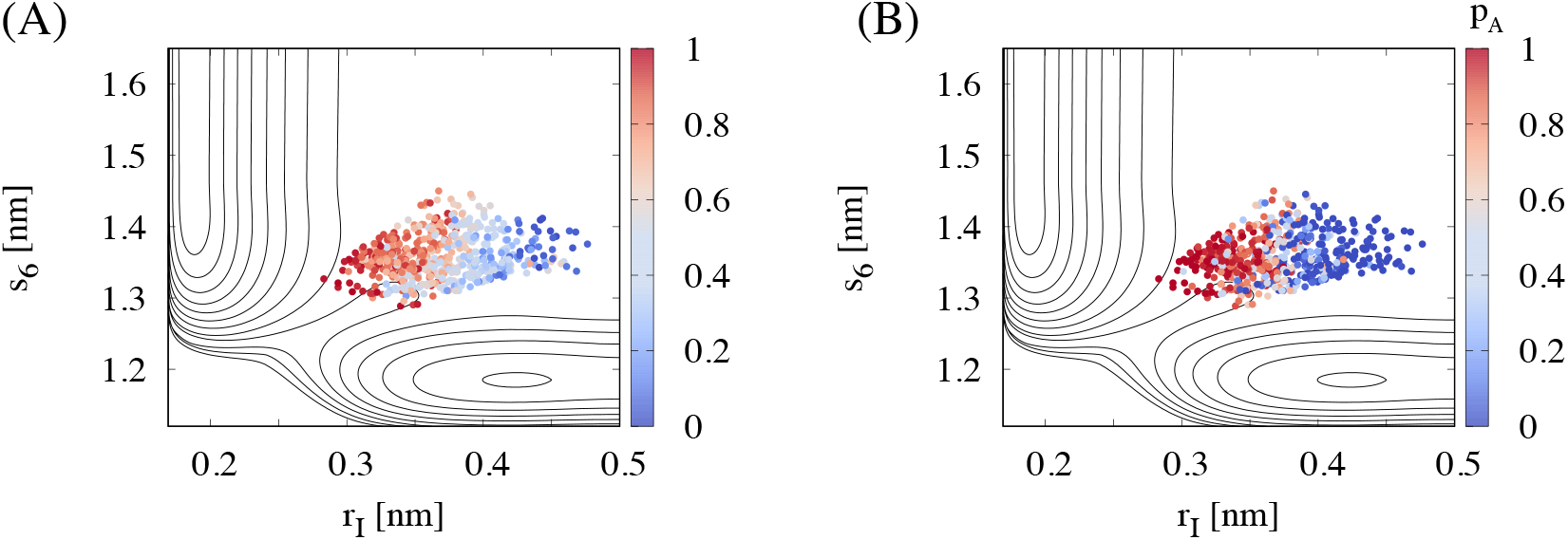
Committor probability for 600 shooting points as function of the features *r*_*I*_ and *s*_6_. (A) Committor probability for conformations from transition path sampling. (B) Committor probability after a constrained equilibration in which the positions of Mg^2+^, RNA, five closest water molecules and exchanging water were kept constant and the solvent was relaxed. The energy contour spacing is 5 *k*_B_T.

The Steinhardt-Nelson order parameters measure to which degree the hydration shell is ordered. The 3rd Steinhardt-Nelson order parameter around atom *i* is calculated from

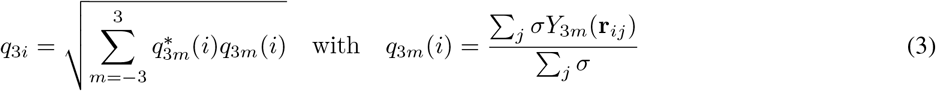

where *Y*_3*m*_ is the 3rd order spherical harmonics and *σ* is a switching function. *σ* is one for the atoms of the hydration shell under consideration and zero otherwise.

The 4th Steinhardt-Nelson order parameter is calculated from

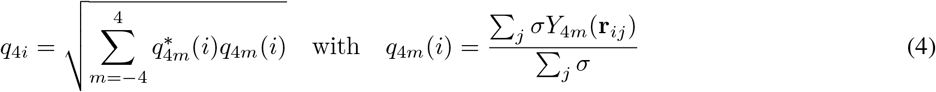

where *Y*_4*m*_ is the fourth order spherical harmonics.

The 6th Steinhardt-Nelson order parameter is calculated from

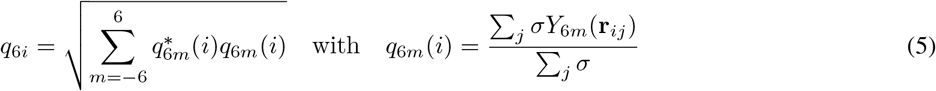

where *Y*_6*m*_ is the sixth order spherical harmonics. All distances, angles, Steinhardt-Nelson order parameters and tetrahedral order parameter were calculated with PLUMED. The number of hydrogen bonds were calculated with GROMACS.

**Table S1:**
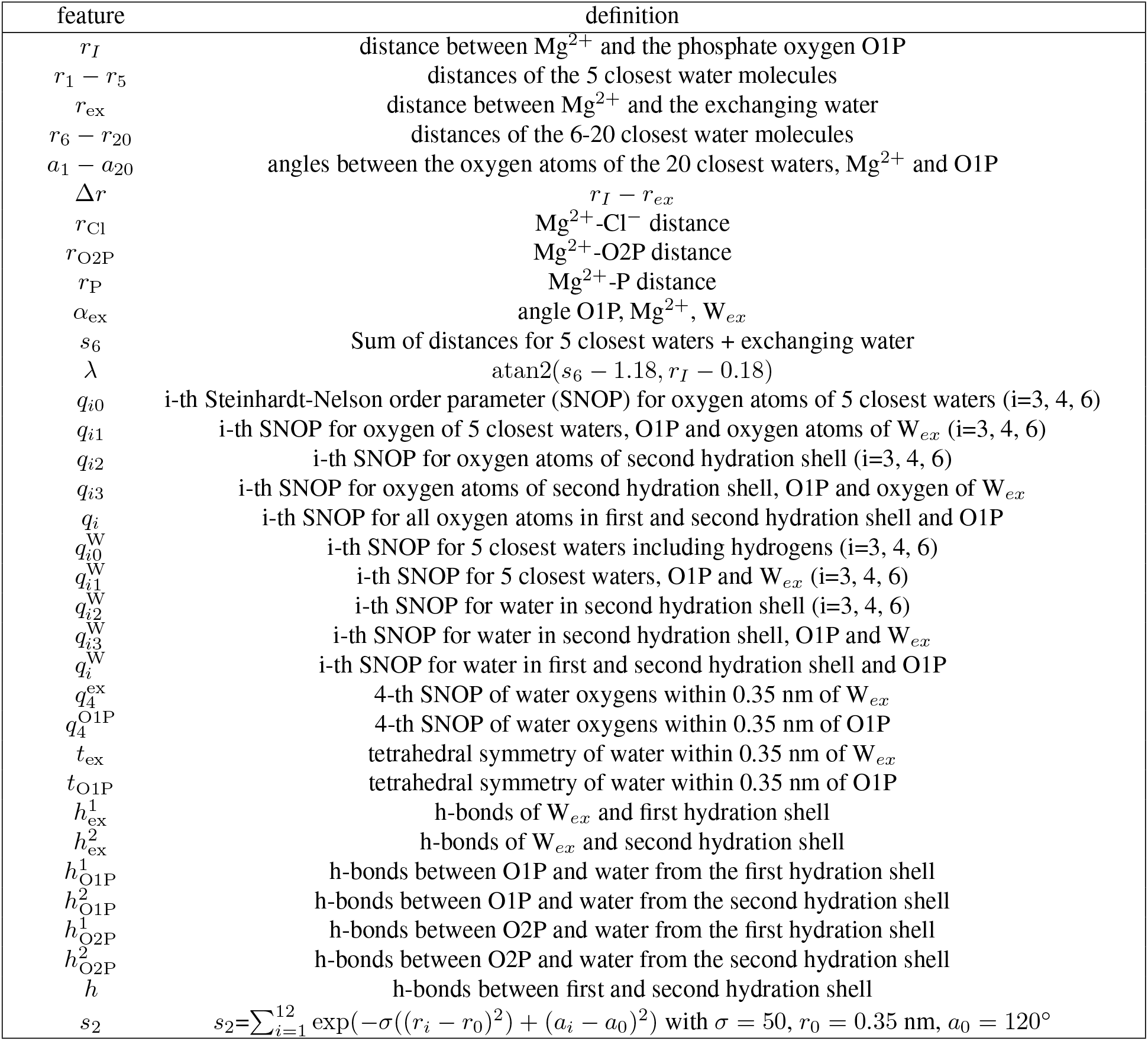
Features used for deep learning.

**Table S2:**
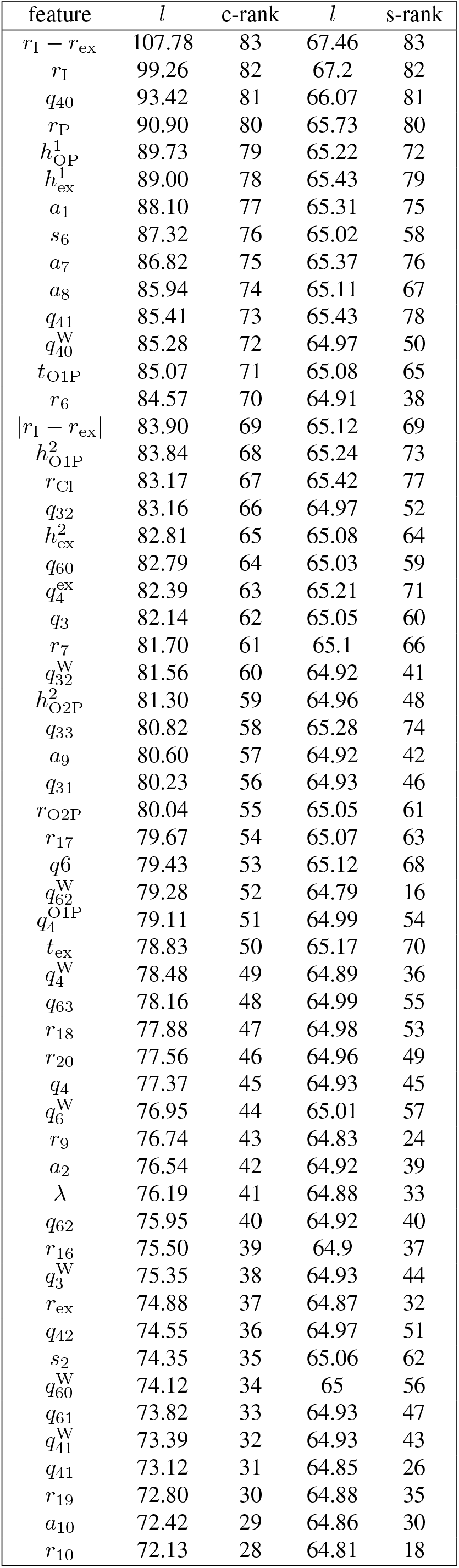
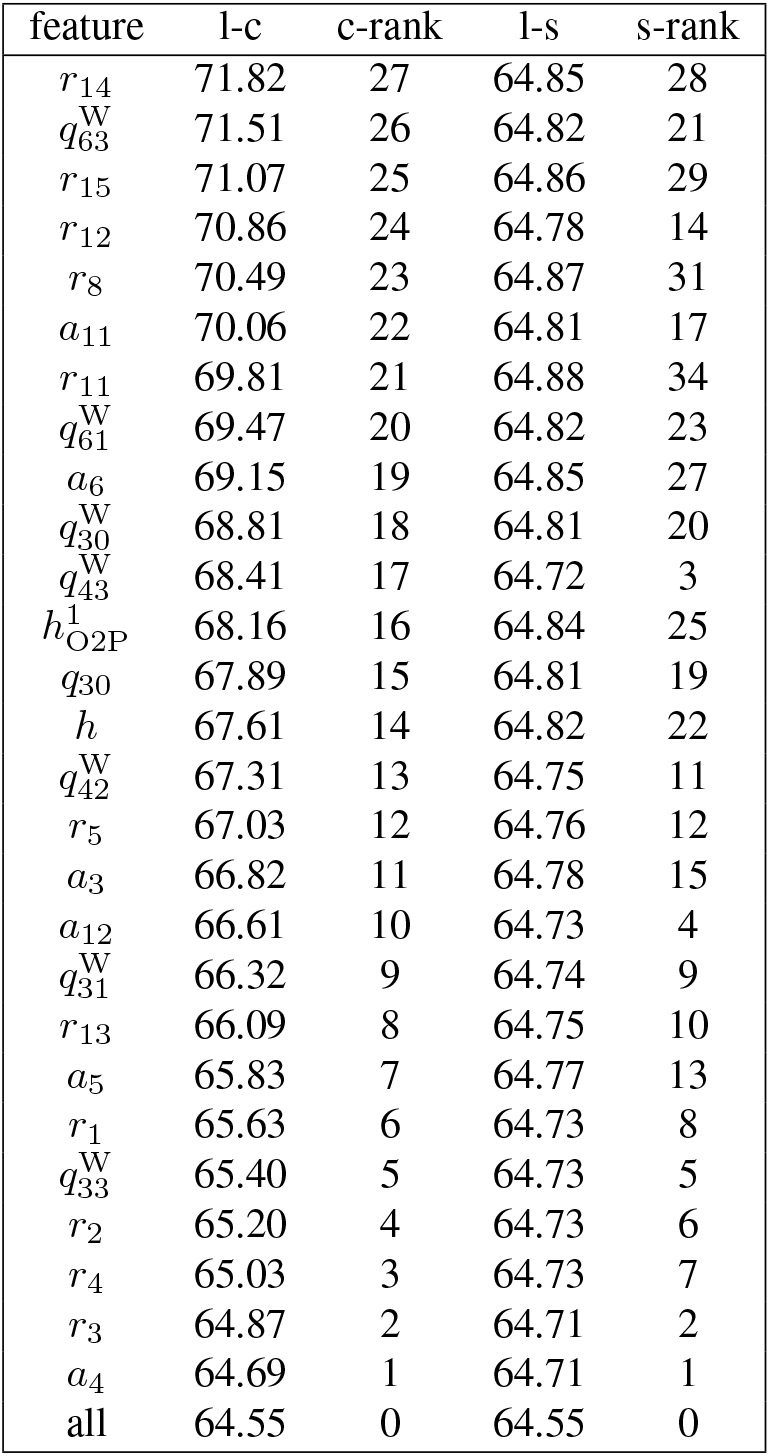
Feature ranking according to combined and single feature relevance and negative log likelihood loss *l*.

## Notes

### Competing Interest Statement

The authors have declared no competing interest.

